# Microbiology and Anthropology: a multidisciplinary approach for estimating time since death during Winter in rural New South Wales, Australia

**DOI:** 10.1101/2024.11.05.622197

**Authors:** Hayley Green, Bjorn Espedido, Rogine Ligot, Slade O. Jensen, Charles Oliver Morton

## Abstract

Accurate determination of the post-mortem interval (PMI) is critical in forensic casework. Most studies conclude that PMI determinations are dependent on local climatic and geographical factors. Despite this, there is little data in an Australian environment outside of entomological studies. In the absence of insect data on or around the remains, alternative methods are required.

Anthropological methods of observing and scoring the extent of decomposition at the time of discovery provide only broad estimates of PMI. Microbial succession is responsible for these observable soft tissue changes, particularly in early and late decomposition. The aim of this study was to combine anthropological and microbiological methods to provide data for determining PMI in a temperate Australian climate.

Microbial DNA was isolated from skin and cavity swabs and used to perform 16S rRNA metagenomic analysis of pooled DNA samples to allow the choice of target taxa for qPCR. qPCR indicated significant changes in microbial communities with a dominant population of *Gammaproteobacteria* at early time points giving way to *Firmicutes* and *Bacteroidetes* near the end of the experiment. Expressing qPCR data as a ratio of *Lactobacillales*/*Enterobacteriaceae* provided data that could be useful in determining early and late decomposition. The genus *Psychrobacter* was identified as a good indicator of late decomposition in winter conditions. qPCR analysis, with further refinements, could be part of an accurate quantitative method of determining PMI.

## Introduction

Through popular television programs such as the “CSI: Crime Scene Investigation” franchise, the general public has developed a fascination with and respect for forensic science. This has highlighted the importance of forensic science research and its significance as part of the criminal justice process (1).

Forensic science is concerned with collecting, identifying, and interpreting physical evidence such as fingerprints, bloodstains, hair, soil, and DNA. One of the most difficult types of physical evidence to identify is the amount of time that has lapsed since death, also known as the post-mortem interval (PMI). Accurate determination of PMI is critical to every death investigation as it facilitates identification of the victim and suspects by reducing the potential pool of missing persons to which the remains could belong and confirming or discrediting witness testimonies should the case be heard in a court of law (2, 3). Difficulties in determining an accurate PMI can be attributed to the fact that there is a poor understanding of the process of decomposition of a body after death (2). Decomposition is a complex process, driven by many factors such as biochemical reactions, insect activity and bacterial activity (4). Adding to the complexity is that each of these factors is dependent on geography and climate (3, 5, 6).

In instances where a forensic anthropologist is required to estimate PMI, the most common method used is the determination of the most advanced ‘stage’ of decomposition present on the deceased at the time of discovery and correlating the observations to the anthropologists knowledge of the environmental region in which the remains were found (3). This method of estimating PMI is qualitative and typically produces broad estimates measured in months or even years (3).

Recently, forensic anthropologists have started to explore ways to develop more quantitative and less subjective methods. These methods typically involve linking the qualitative soft tissue changes observed on decomposed remains at the time of discovery to a measure of temperature (3) and/or other taphonomic factors such as humidity and whether remains decomposed in aerobic or anaerobic conditions (7). These methods, however, were developed internationally and, as it is widely recognised that decomposition processes are climate and environment dependent (3, 5, 6), may not be applicable in Australian conditions (8).

Despite the semi-quantitative nature of the Megyesi et al. (3) and Vass (7) methods, quantitative methods have been found to only be effective during the early stages of decomposition, when the expertise of anthropologist is less likely to be required (9). These methods include measuring vitreous potassium concentrations in the eye (10), insect succession (11) and decomposition of body organs (12), which provide relatively accurate PMI determinations that range between hours during the first two weeks post-mortem (12). Vass (7) also determined that the ‘universal’ anthropological method developed in his study could only be used during the pre-skeletonisation phase of decomposition, and that it would not be effective when the remains were mummified or skeletonised. This is problematic, as anthropologists are typically required to determine PMI with long post-mortem intervals (9). A method that assists anthropologists with their PMI determinations during advanced and dry stages of decomposition is thus warranted.

Recently, a novel approach of estimating time since death that correlates morphological changes to soft tissue and bone after death with numbers and types of microbes present during decomposition has recently emerged in the literature (2, 13, 14). They can be considered a ubiquitous form of physical evidence due to the stability and predictability of microbes in the environment (15). Results indicate that bacterial communities have a vital role in the process of decomposition, with communities from both the host and the associated environment changing in a specific, reproducible way (16). Research has focused on both animal and human models, examining microbial community changes from samples sites on the host organism such as the skin, rectum, cavities of the face (eyes, ears and nose), internal organs and nose in both controlled lab experiments and outdoor experimental scenarios (17) as well as forensic cases (14). Despite these studies, the most appropriate sample site for estimating PMI using microbes is still yet to reach a consensus. Analysis of the microbiome on tissue in more advanced stages of decomposition, such as skeletonised remains, has demonstrated that PMI estimation is possible (18), but more studies are required. The results of previous studies suggest that bacterial community changes over time during decomposition could be explaining the visual changes that anthropologist rely on to make their PMI estimations. It also suggests that measures of the microbiome over time may be beneficial in estimating PMI in the more advanced/dry stages of decomposition when existing PMI methods provide estimates of years at best. Major limitations of these microbial studies are that they were conducted internationally, and that microbial activity may be geographically and seasonally specific particularly for microbes in the soil (19). To date, there are no Australian specific studies published.

Many of these studies have employed high throughput sequencing methods to examine microbial populations (13, 16). Such studies have also revealed the potential of microbiome date to assist in the determination of PMI (20, 21). This approach gives a wealth of information about microbial communities but requires specialist skills to analyse and interpret. Quantitative polymerase chain reaction (qPCR) can be used for a variety of purposes from diagnosis of infectious disease (22), gastric medical conditions (23), to monitoring microbial communities (24, 25). qPCR has also been employed in testing of gut microbial communities to determine PMI (26). qPCR methods can give same day results that are specific, broadly available, and easy to interpret. This gives an alternative DNA-based approach to sequence analysis that can assist in the determination of PMI.

The aim of this study is therefore to determine baseline data on the change to microbial communities during decomposition in an Australian climate for the first time. Specifically, community diversity will be assessed during each stage of decomposition (3, 6, 7, 27–29) to determine if changes to the microbiome correlate to the gross anatomical changes observed at each time point. We hypothesise that bacterial community diversity will change dramatically and measurable over time, providing a useful tool that anthropologists can utilise in PMI estimations, particularly during advanced stages of decomposition.

## Materials and Methods

The study was carried out over a 12-week period during the 2014 season of winter (June 10^th^ - September 10^th^) at Western Sydney University’s Hawkesbury campus in Richmond, NSW (GPS coordinates: 33.61° S, 150.75° E). The location of the research site on Hawkesbury campus is an area comprised of dense bushland with an abundance of *Melaleuca decora* (paperbark) trees (Figure 1a). The research site is isolated from the general public but periodically, cattle and sheep were given access to the field site for grazing. Wildlife such as kangaroos and foxes were also observed at the site. The Hawkesbury region experiences mild winter temperatures, with an average daily temperature of 14°C and a low below 5°C with very little rain and no snow (CSIRO, 2007).

**Figure 1.**
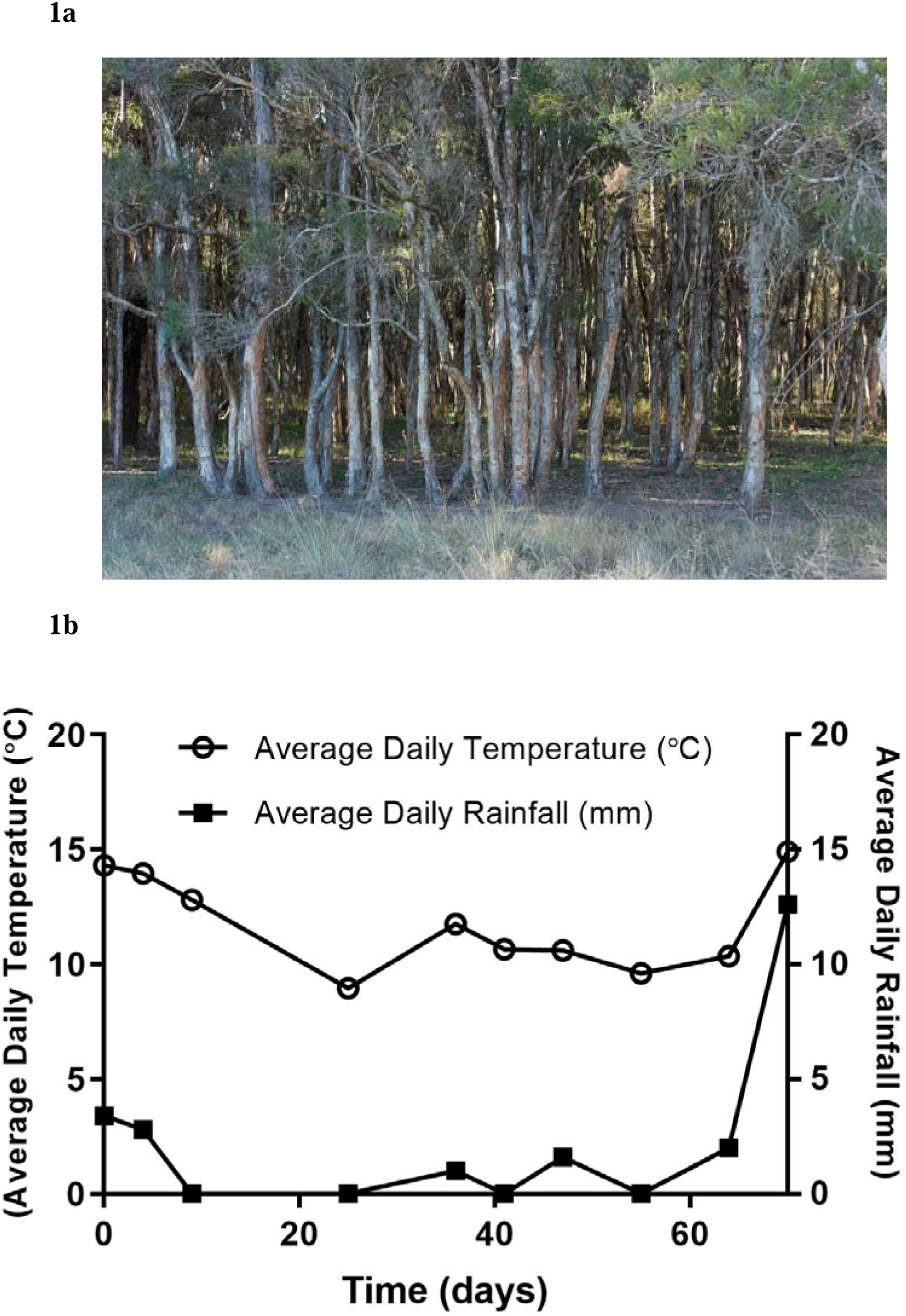
(**a**) Research site comprising of a dense *Melaleuca decora* (paperbark) tree forest. (**b**) Climate data for the experimental period. The average daily temperature (°C) and the average daily rainfall (mm) were acquired using dataloggers and plotted for the duration of the experiment (70 days).

Four adult pig carcasses (*Sus scrofa domesticus*) weighing approximately 70 kg each, were obtained post-mortem from a local abattoir. Carcasses were killed by a captive bolt to the brain and were immediately transported to the research site via an enclosed trailer. The study site is located approximately 20.9 km or 22 minutes from the abattoir. Within an hour after death, the carcasses were placed on their left side on the natural soil-based surface of the ground. This was recorded as Day 0. Remains were contained in metal cages to prevent vertebrate scavenging but were exposed to normal weather conditions and invertebrates still had access to the remains.

Visual observations of the stage of decomposition of each of the carcasses were recorded and photographed, and bacterial samples from the eye, ear, nostrils, anus, and abdominal skin (Figure 2a) were collected twice a week during the 12-week study period. Gross anatomical changes during decomposition were described according to Galloway et al. (27) and Megyesi et al. (3). Temperature data were collected using onsite Tiny Tag Plus 2 dataloggers attached to one of the cages, while humidity and rainfall was taken from freely available data from the nearest weather station in Richmond, NSW via the website of the Bureau of Meteorology (BoM).

**Figure 2.**
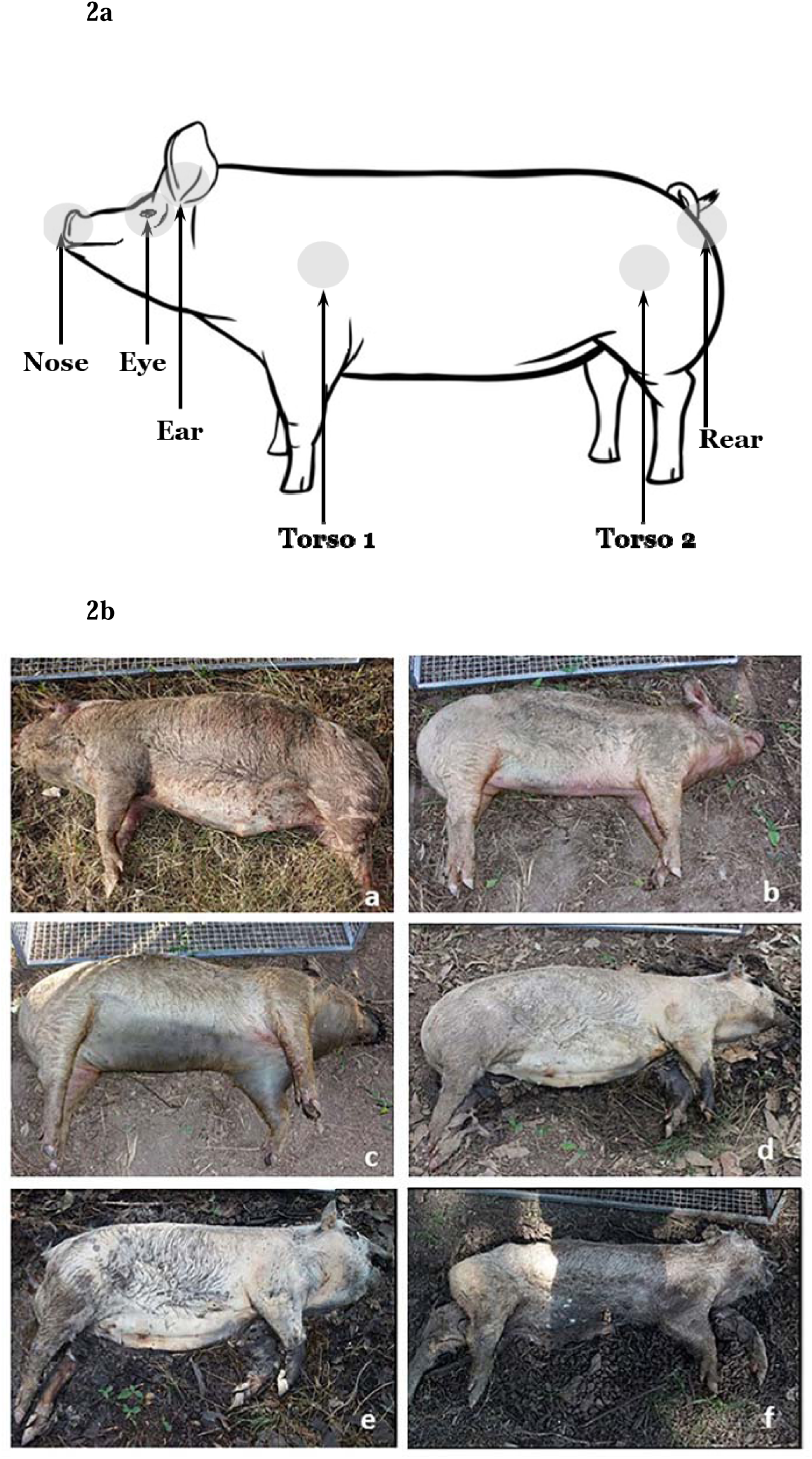
(**a**) Locations of sampling sites on the body of the pig for bacterial population analysis. Each site was swabbed periodically to collect bacteria; the swabs were stored at - 20°C after sampling. Representative summary of the average morphological changes observed over the 12-week project duration. (**b**) Representative photos of the stages of decomposition. a) ‘fresh’ stage with evidence of lividity and marbling; b) ‘early decomposition’ characterised by green discolouration of the abdomen; c) end-stage ‘early decomposition’ with extensive discolouration of the abdomen; d) ‘advanced’ decomposition with skin slippage of the limbs and bone some exposure of the facial bones; e) biomass loss after a period of rainfall; f) drying of tissues around Day 70. No further soft tissue changes were observed between this time point and the completion of the study (Day 90).

Bacterial samples were collected using sterile cotton buds from six sites on the animal: ear canal, eye, nostril/nasal cavity, rectum and skin of the shoulder and flank. These were immediately placed in a 1.5 ml Eppendorf tubes and stored at -20°C until DNA extraction, which began 12 weeks after Day 0.

### Isolation of bacterial DNA from swabs

Bacterial DNA was isolated from swabs using the Isolate II Genomic DNA Kit (Bioline) according to the manufacturer’s instruction with some modifications. Briefly, frozen swabs were allowed to thaw at room temperature for 15 min, and then incubated in a 1.5 ml microcentrifuge tube containing 180 µl of buffer GL with 25 µl proteinase K solution at 56°C for 120 min, tubes were vortex mixed for 15s every 30 min. The swab was removed while pressing against the wall of the tube to retain lysate, 200 µl of buffer G3 was added and the mixture incubated at 70°C for 10 min. Finally, 210 µl ethanol (96-100%) was added; the sample was mixed and added to a DNA binding column. The manufacturer’s instructions were then followed without deviation and the sample was eluted in 100 µl of elution buffer.

### Bacterial 16S amplicon sequencing pilot study

To determine the overall composition of the bacterial community on the experimental pigs throughout the course of the experiment 16S rRNA gene amplicon sequencing was used to survey the bacterial population on the pig carcasses. Four time points were chosen (0, 9, 40 and 70 days) that corresponded to major changes in decomposition. DNA samples at each time point were pooled and bacterial 16S rRNA gene fragment libraries were prepared using the Ion 16S Metagenomics Kit (Thermo Scientific) and sequenced using the Ion Torrent Personal Genome Machine (PGM; Thermo Scientific); as part of the metagenomics kit, 7 of the 9 16S rRNA gene hypervariable regions were amplified for sequencing (V2, V3, V4, V6, V7, V8 and V9). On completion of the sequencing run, data was automatically uploaded to the Ion Reporter^™^ Software (Thermo Scientific) for quality control, read mapping, annotation, and reporting. As part of this process, reads were compared to two 16S rRNA gene databases (MicroSEQ^®^ ID database, Thermo Scientific and the Greengenes database, Lawrence Berkeley National Laboratory) for phylogenetic assignment and metagenomic analyses.

### qPCR of bacterial populations

Bacterial taxa for qPCR were then chosen based on changes in abundance over the four time points determined by the 16S amplicon sequencing pilot data (Figure 3). The following published PCR primers were chosen to detect microbial taxa in each body site at each time point: Pan-bacterial (Eub338/Eub518) (30); Phyla-specific primers, *Actinobacteria* (Actino235/Eub518) (30), *Firmicutes* (Lgc353/Eub518) (30), *Bacteroidetes* (Cfb319//Eub518) (30); Class-specific primers, *Gammaproteobacteria* (Gamma887F/Gamma1066R) (31), *Bacilli* (BLS342F/1392R) (32); Order-specific primers *Clostridiales* (16SClost_F/16SClost_R) (33), *Lactobacillales* (F-lac/R-Lac) (34); Family-specific primers, *Enterobacteriaceae* (Uni515F/Ent826R) (35), Genus-specific primers, *Psychrobacter* (432F/476R) (36)

**Figure 3.**
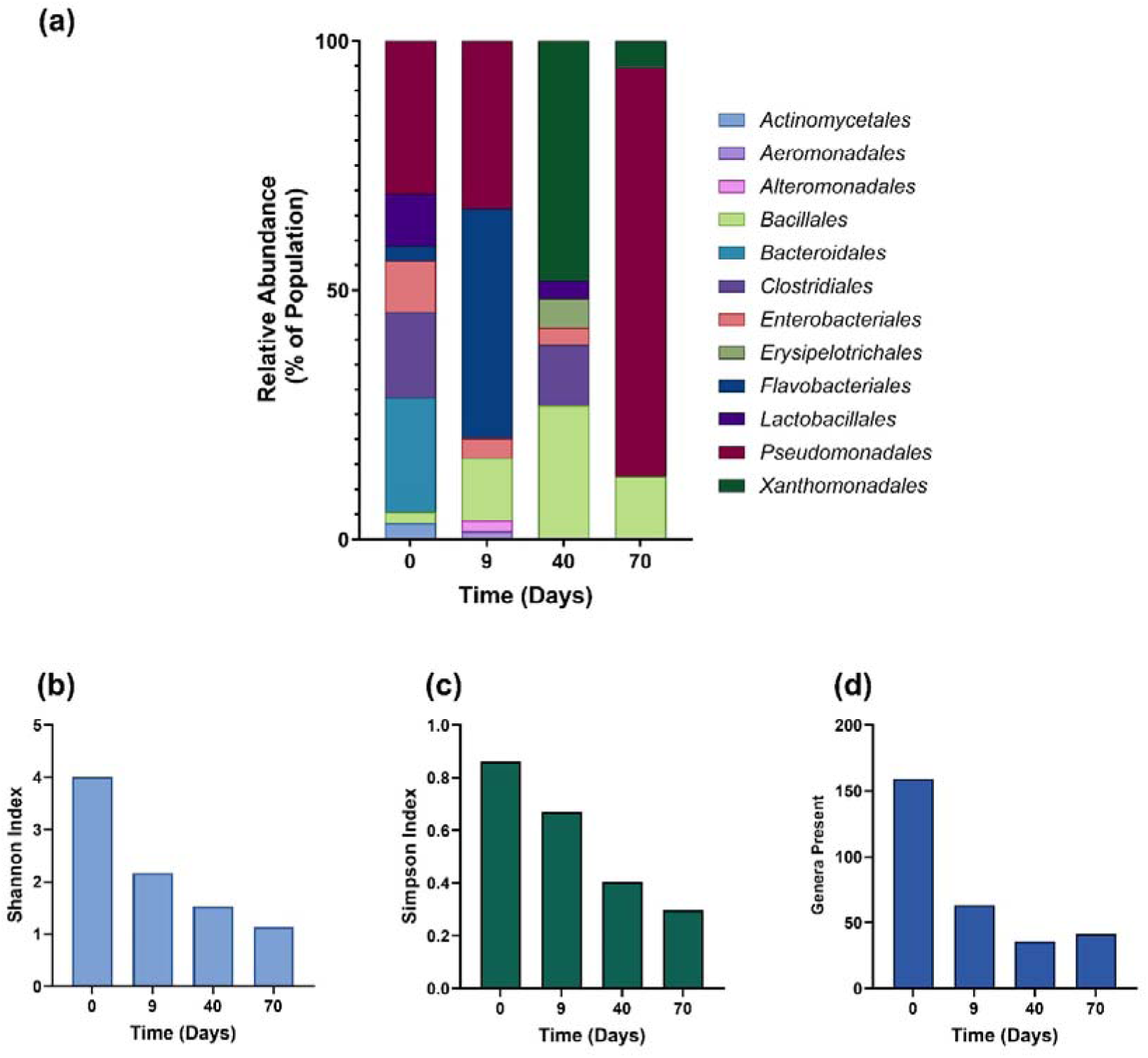
**(a)** Metagenomic data for pooled pig samples for fours timepoints indicative of major stages of decomposition. The charts were generated from sequence analysis of 16S PCR products amplified from pooled DNA samples taken from different body sites at each time point. Colour codes vary depending on the presence of taxa at each time point. **(b – d)** Indicators of species diversity show that there is a decrease in overall numbers of genera present and in species diversity over the course of the experiment.

Bacterial DNA samples were diluted to 10 ng/µl for qPCR analysis. Each 20 µl qPCR assay contained; 10 µl of QuantiFast SYBR Green PCR Kit mastermix (Qiagen), 0.2 µM each primer, 2 µl template (10 ng/µl), and 6 µl PCR grade water (Sigma-Aldrich). The reactions were performed on a 7500 Fast real-time PCR machine (Thermo Fisher) using the following cycling parameters: 95°C for 10 mins initial denaturation then 40 cycles of 95°C for 15s, 50°C for 30s, and 72°C for 30s. This was followed by the instrument’s default melt curve for analysis of SYBR Green qPCR. The Cq values were converted into relative quantity for analysis using the equation 2^(40-Cq)^ (37, 38).

The mean relative abundance for each bacterial taxon at each time point were plotted for all body sites as heat maps. These data were analysed non-parametrically by Kruskal-Wallis and Dunn’s multiple comparison test to analyse differences in bacterial population at each time point using GraphPad Prism version 7. Using this analysis, the assay for *Enterobacteriaceae* was found to be relatively stable over the course of the experiment, so it was chosen as a standard. The assay for *Lactobacillales* showed an increase in relative abundance over time. The abundance of *Lactobacillales* relative to *Enterobacteriaceae* was calculated using the equation 2^-ΔCq^. These results are an example of how a qPCR assay can be used to monitor bacterial abundance and potentially contribute to PMI determination.

## Results

### Climate data

The average daily temperature collected from the dataloggers at the site ranged from approximately 7.7°C -16.2°C (mean= 11.69°C) over the 90 days of the trial. Rainfall was intermittent throughout the trial, documented as occurring only 27/90 days and totalling 125.8 mm. The average daily humidity was 63.33%. The climatic data for the experimental period is shown in Figure 1b.

### Anthropology

All four carcasses remained in the ‘fresh’ stage of decomposition for approximately 6 days (Figure 2b). This stage was characterised by lividity and marbling on the abdomen and limbs and minimal insect presence (Figure 2b, panel a). By post-mortem Day 6, remains exhibited a green discolouration in the lower right quadrant of the abdomen, which was well established by Day 9, signalling the beginning of ‘early’ decomposition (Figure 2b, panels a-b). Early decomposition was characterised by bloating of the remains by day 16 and darkening of the green discolouration, which was visible along the entire length of the carcass by Day 16 (Figure 2b, panel c). By day 21 there was observable maggot activity around the creases of the hind limbs, as well as some purging of fluid from the nasal cavity. However, remains were not classified as being in a state of ‘advanced’ decomposition until between Days 30-41 post-mortem. Remains during this stage exhibited moist decomposition on the areas of the carcasses in contact with the grounds surface which was accompanied by extensive maggot activity. There was also bone exposure in the head region, specifically at the jaw and orbits. Skin slippage was observed on the limbs only, and caving or sagging of the remains did not occur until around day-40 (Figure 2b, panel d). A period of heavy rain occurring between days 68-69 rehydrated the tissues and resulted in a period of further biomass loss (Figure 2b, panel e) and internal active decay. Drying of tissues (mummification) was achieved by Day 70 (Figure 2b, panel f), with minimal further physical changes observed between day 70 and the conclusion of the trial.

Based on the visual observations by the anthropologist, it was determined that the bacterial samples from days 0, 9, 40 and 70 would be subjected to 16S metagenomic analysis, as these were the time points when all pig carcasses exhibited typical gross anatomical changes associated with each decomposition stage as defined by Megyesi et al. (3) and Galloway et al. (22). The accumulated degree day (ADD) at each sampling point was also calculated as a point of comparison with previous studies, however, research has indicated that ADD as a predictor of PMI in Australian contexts is inaccurate (24).

### Microbial Analyses

An initial metagenomic analysis of pooled samples at days 0, 9, 40 and 70 indicated changes in the bacterial population structure over time (Figure 3). This pilot analysis revealed that bacterial diversity decreased over time (Figure 3 a – d). These sequence data provided the basis for the selection of oligonucleotide primers for qPCR analysis of each body site samples at each time point.

Further analysis of all samples at all time points using 16S qPCR for selected high-level taxa showed changes in the abundance of the *Gammaproteobacteria*, Firmicutes, *Bacteroides*, and *Actinobacteria* over the course of the experiment (Figure 4). All sample sites (Figure 4 b – g) showed an increase in the proportion of *Bacteroidetes* and *Firmicutes* over time with a relatively lower abundance of *Gammaproteobacteria*. This can be linked to the overall abundance of bacteria (Figure 5a), as the population density decreased the abundance of some taxa increased relative to the remaining population. These analyses differ to the metagenome (Fig 4a) but that was a pooled sample analysed by a different methodology.

**Figure 4.**
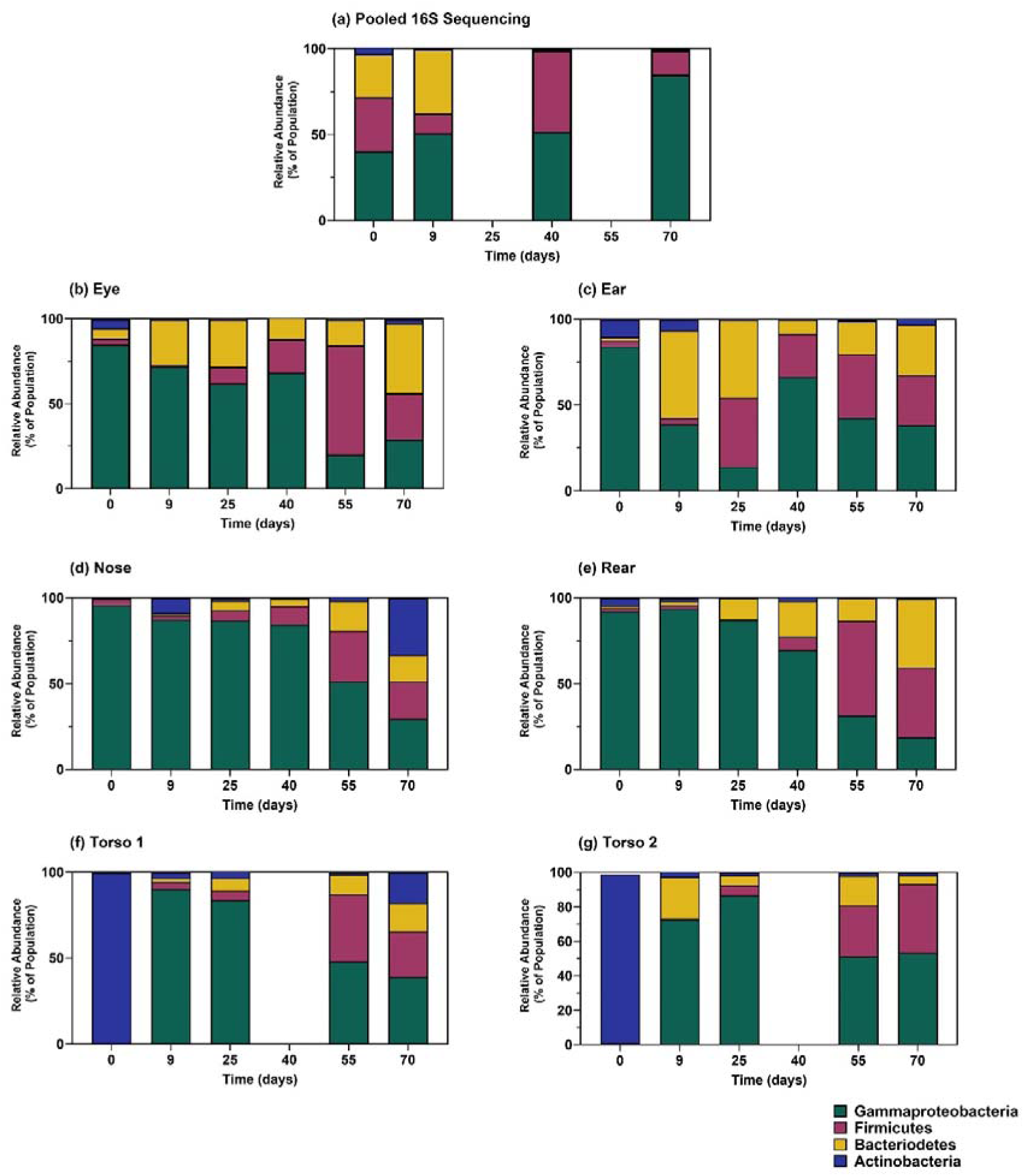
qPCR data for the abundance of higher taxonomic classes for each sampled body part at each sample time. The Metagenomic data was limited due to cost, so qPCR was used to monitor the abundance the different groups of bacteria (*Gammaproteobacteria*, *Firmicutes*, *Bacteroidetes*, and *Actinobacteria*). These data represent mean percentage abundance (to match the presentation of the metagenomic data); from three pigs. There was a similar trend for each sample site with *Gammaoproteobacteria* comprising a smaller proportion of the overall population as time progressed. Data presented are means from three replicate animals.

**Figure 5.**
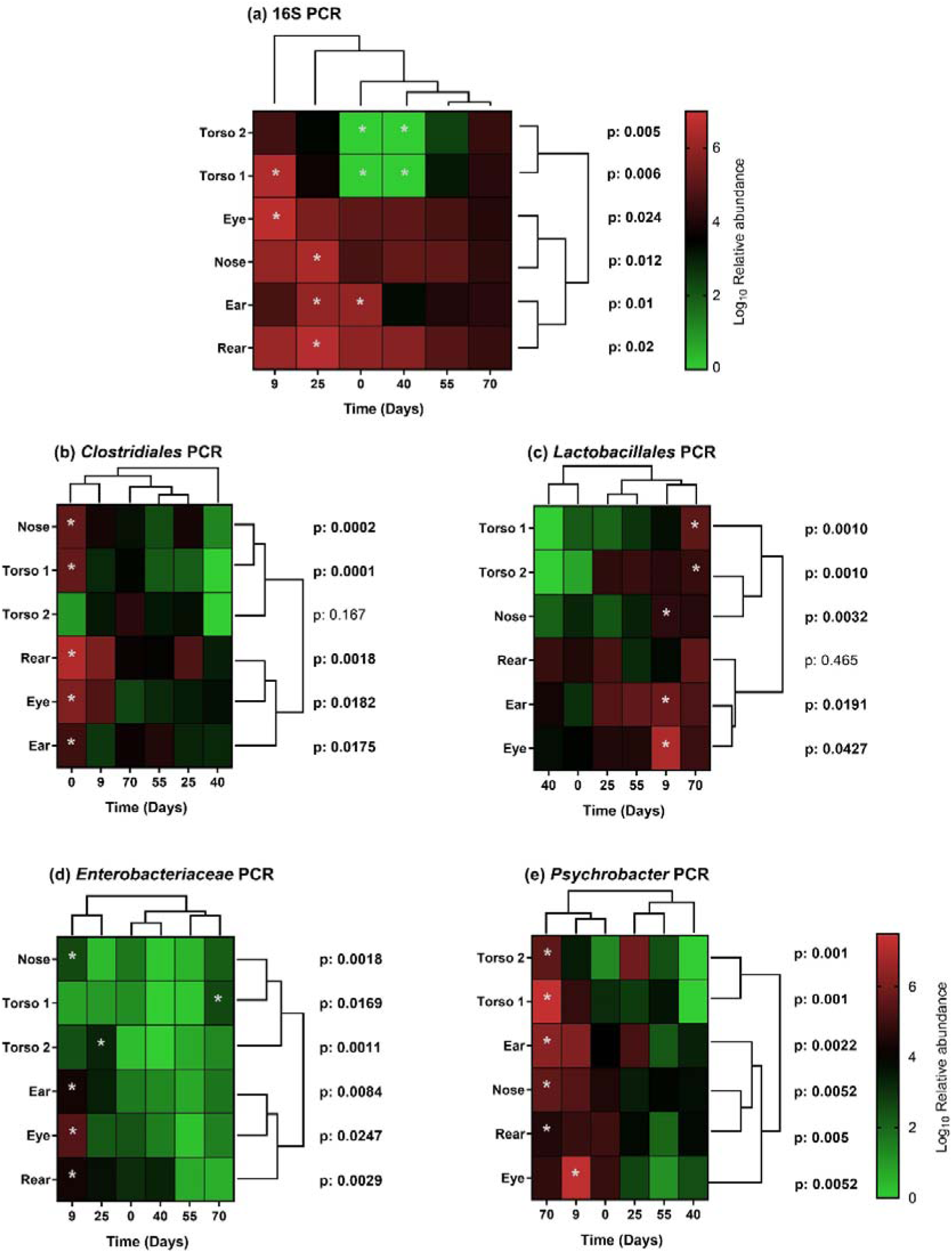
Heat maps showing temporal changes in bacterial abundance during the experiment. (a) Bacterial abundance measured by 16S qPCR. Data presented as mean relative abundance determined for three pig carcasses. (b) Changes in the abundance of specific bacterial taxa. The heat maps represent relative quantities of bacteria determined by qPCR using primers specific for the *Clostridiales*, *Lactobacillales*, *Enterobacteriaceae*, and the genus *Psychrobacter*. Data presented are as log_10_ of the relative abundance value (mean of data from three pig carcasses). Data were analysed in each sampling site across each sampling time point by Kruskal-Wallis testing with Dunn’s multiple comparisons (* p < 0.05).

### Bacterial abundance and climatic factors

16S qPCR indicated that the greatest abundance of bacteria was observed at earlier time points (Days 0, 9, and 25), with the lowest abundance at 70-days when the carcasses showed extensive decomposition (Figure 5a). The torso T1 site had the least bacterial DNA whereas the eye had the greatest (at day 9), however after 40 and 70 days the amount of bacterial DNA was similar for all body sites and almost 100-fold lower in abundance than at earlier time points. Bacterial DNA could not be quantified at the 40-day time point for the two torso sample sites, this may have been due to climatic factors. The torso sample sites appeared to track the drop in rainfall (Figure 1b). These exposed sites are more directly affected by the climate.

### Dynamics of bacterial taxa during decomposition

Taxa specific qPCR was performed to see if qPCR could be used to monitor bacterial abundance and potentially determine PMI as a cheaper and speedier alternative to metagenome sequencing. The qPCR data was presented as heat maps showing changes over time since death at each body site (Figure 5).

Analysis of the specific taxa provided better discrimination between time points than the pilot metagenome analysis, so qPCR was performed on *Lactobacillales*, *Clostridiales*, *Enterobacteriaceae*, and *Psychrobacter* at each time point for each body site (Figure 5 b - e). The qPCR data for these was highly variable, although there were statistically significant results associated with single time points for the taxa tested. Variability is inherent when examining microbial populations, there will often be differences in absolute abundance between samples. To account for this variability required a reference taxon, in this case *Enterobacteriaceae* was chosen as the reference, since it had a relatively stable abundance over time (Figure 5d). The abundance of *Lactobacillales* increased over time (Figure 5c) so the ratio of *Lactobacillales*/*Enterobacteriaceae* (L/E ratio) was used to plot changes in bacterial abundance over time (Figure 6). These data show an increasing L/E ratio over time that was consistent between body sites.

**Figure 6.**
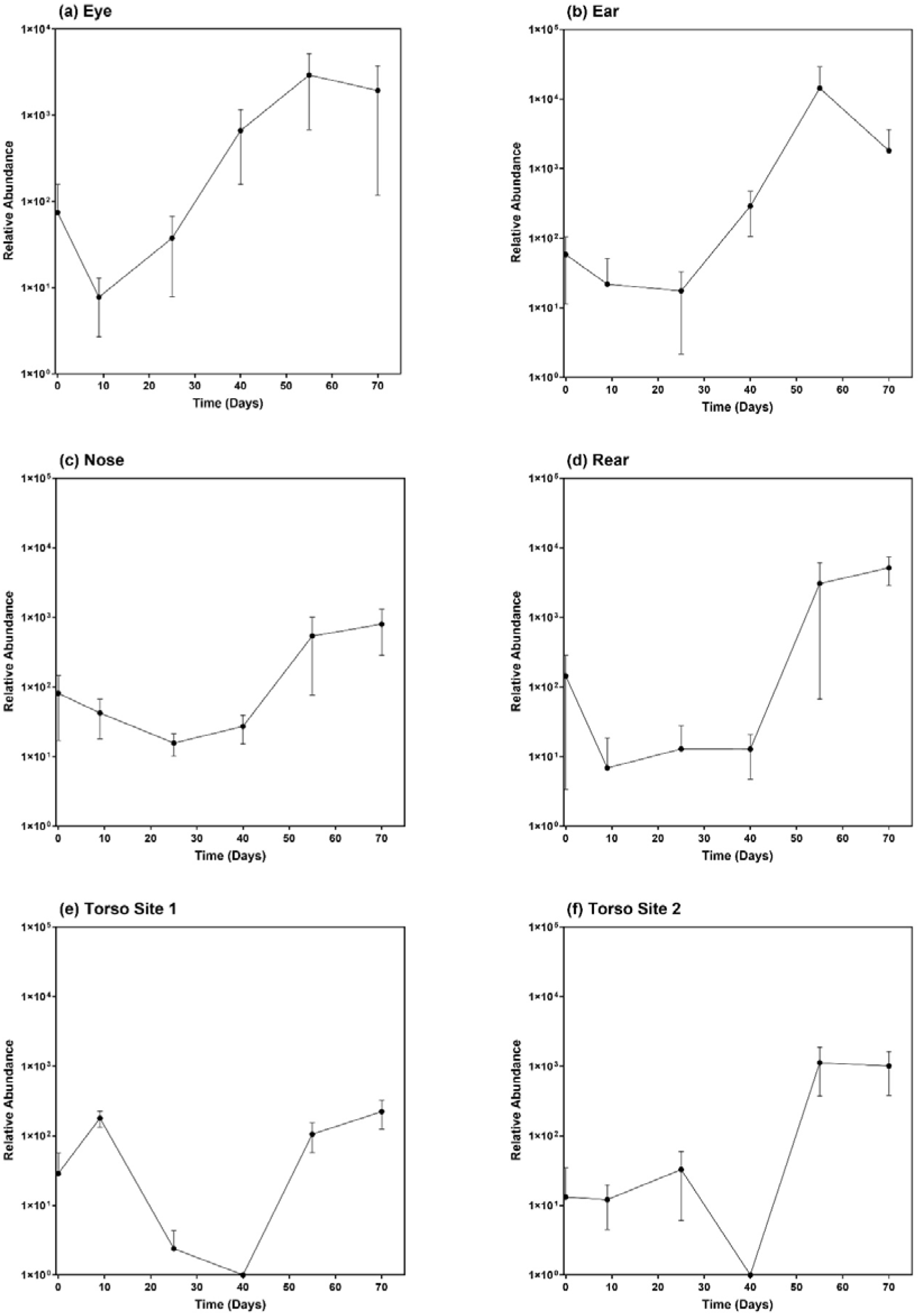
Tracing the relative abundance of *Lactobacillales* relative to a reference taxon (*Enterobacteriaceae*) over the course of the experiment. *Lactobacillales* were expressed relative to *Enterobactericeae* using the equation 2^-ΔCq^. The raw qPCR data per species is not easily interpreted but when the ratio of *Lactobacillales*/*Enterobacteriaceae* is considered then the interpretation is clearer. In the eye, ear, rear, and second torso site the relative amount of *Lactobacillales* increased across the experimental period. This gives a data point that could help to determine for early and late decomposition. Data are mean and standard error from three replicate animals.

The L/E ratio could assist in determination of PMI based, based on these data it was most consistent for the eye, ear, and rear sampling sites; the surface sites were more variable and there was the possibility of no sample recovery which occurred for the 40-day samples.

## Discussion

The present study is an important contribution to multidisciplinary approaches to studying decomposition processes in a temperate climate. The use of an animal model in lieu of available human cadavers enabled for the control of the body condition, which is critical for establishing whether the succession of microbes during decomposition is predictable (14, 39). A winter season was chosen for this study so as anthropological results could be comparable to previous studies conducted in similar environmental contexts (16, 40, 41) however the authors acknowledge that the Australian climate is not necessarily directly comparable to northern hemisphere studies. Additionally, while seasonality has been recognised as having an effect on both the progression of remains through stages of decomposition and the composition of microbial communities, winter studies are largely under-represented in the literature (41). By focussing on variability in decomposition and the post-mortem microbiome in a cold climate, the study was able to capitalise on longer data collection periods due to slow decomposition rates and provide baseline post-mortem microbiological data in an Australian context for the first time.

The timing of the progression of decomposition through the ‘fresh’ (day 6 - 9), ‘early’ (day 9 - 30) and ‘advanced’ (day 31 - 41) stages and the morphological changes observed during each stage were consistent with previous studies (16, 42). This ‘typical’ pattern clearly defined sample collection timepoints for microbiome analysis at each decomposition stage (Day 0, 9, 40 and 70). The final time point (Day 70) was chosen to determine if the increase in decomposition activity after a period of rainfall (Day 68 - 69) had an effect on the abundance and composition of the microbiome.

The microbial element of this study initially consisted of 16S metagenome analysis of pooled samples (different body sites and experimental animals) at four time points to identify key taxa in the microbial population that were changing over time (Figure 3a). This revealed that microbial diversity was greatest early in decomposition with a pronounced decrease in species diversity by the end of the experimental period when soft tissues had begun to desiccate (Figure 3b-d), with these results in agreement with other studies of decomposing pigs (13). qPCR targeting 16S revealed a decrease in the overall quantity of bacterial target DNA over the course of the experiment (Figure 5a), previous studies using qPCR have also shown a gradual reduction in bacterial abundance over time (26).

Amplicon sequencing analysis of each body site at each time point was not financially viable for this study and qPCR of selected taxa (from our metagenome analysis) was selected as the best alternative. This may be the case for many centres. Analysis of each body site at each time point showed variability in the major bacterial taxa, *Gammaproteobacteria*, *Firmicutes*, *Bacteroidetes*, and *Actinobacteria*, that were targeted (Figure 4), these taxa also dominate the human microbiome (43). The observed changes in microbial population during decomposition (Figures 4) are consistent with other studies that have measured changes in microbial community structure throughout decomposition (13, 44).

The eye and ear showed similar changes in bacterial diversity which were distinct from the other body sites over the first 40-days, the proportions of bacterial taxa became more similar across all body sites at 55 and 70-days (Figure 4). It is difficult to directly compare these results to previous studies, as non-human mammalian species have predominantly focused on the usefulness of sample sites such as the skin (2, 39, 45) and rectum in estimating the PMI (2), while nasal, eye and ear cavities (46) feature in human studies, as do the previously mentioned sample sites (26, 45, 47). Each site, irrespective of species, has demonstrated promise in estimating the PMI, however, there is yet a consensus on the most useful sample site(s) for PMI estimation (34). The data in this study (Figure 4 f & g) would suggest that the skin may not be the most reliable site for sample recovery.

Studies of the post-mortem microbiomes have indicated that as bodies decompose there is a change in the composition of microbial populations with relatively clearly defined early and late populations. The change in microbial population is thought to be due to the early population being derived from the microbiome of the body and the later population being from the environment (26, 45). The composition of the body changes markedly in terms of oxygen availability, nutrients, and water content which also creates a selection pressure on the composition of the local microbial population. The climatic conditions in the current study caused the carcasses to mummify indicating that there was a loss of moisture through the initial period (0 to 55 days) of decomposition. There was precipitation and a rise in average temperature at around 70-days (Figure 1b) which may have led to the observed recovery in some bacterial populations at the end of the experiment (Figure 5b). It has been reported that seasonal variation in climate affects microbiome analysis in cadavers and the results of this winter study would not be comparable to a study in summer (48, 49).

The ability to contribute to PMI determination requires the identification of characteristic changes in the microbiome over time (14). A qPCR study targeting *Bacteroides* and *Lactobacillus* in the post-mortem human large intestine showed that populations of these bacteria declined over time in a reproducible manner over the first 21 days of decomposition indicating that either genus could be used as an indicator of PMI (26). Measurement of the relative abundance of microbial taxa by qPCR has also been used to monitor gut dysbiosis (50). This article also highlighted that there were no significant differences between results from sequencing and qPCR, adding weight to the use of qPCR for PMI analysis. We chose the L/E ratio to act as a potential indicator of PMI. These data (Figure 6) suggested that this approach could determine if a sample was taken from early or late decomposition. The ratio of *Clostridiales* to *Enterobacteriaceae* was another candidate that we could have chosen for aiding in estimation of PMI. Using the *Clostridiales* is supported by studies of humans where the genus *Clostridium* has been proposed as a key taxon in the determination of PMI. This was true especially in studies of internal organs where the genus was ubiquitous throughout decomposition (51).

The most abundant taxon by the end (Day 70) in our Winter study was *Psychrobacter*, which was a minor component of bacterial diversity at the beginning of the study (Days 0 – 40). The abundance of *Psychrobacter* was significantly greater at 70-days for all body sites except the eye, which reflects the 16S metagenome data (Figure 3a, *Pseudomonadales*). This genus is generally resistant to cold so it should be able to outcompete mesophilic bacteria, particularly those from the antemortem microbiome, during Winter. Bacteria from the genus *Psychrobacter* form part of the antemortem microbiome in pigs and some species are commensals in humans but not to the same extent as in pigs (52, 53). *Psychrobacter* spp. have also been detected at relatively high abundance in Winter grave soils, so they can originate from the surrounding environment rather than just the host microbiome (16). In a study using microbial populations to estimate PMI in cold environments, *Psychobacter* was identified as a key predictor genus for PMI (54). This supports our findings regarding the potential importance of *Psychrobacter* as an important genus at the later stages of decomposition in Winter.

There is a need for linkage of environmental data to microbial populations for microbial population data to be accurately used in forensic analysis (13). qPCR used to monitor bacterial populations can identify early and late decomposition but lacks the greater resolution needed for more subtle changes based on the qPCR targets used in this study (Figure 6). In future studies it would be optimal to use a larger sample size, in different seasons. The DNA isolation method would need to be adapted. The samples became more similar to environmental samples as decomposition progressed, this would require alterations to the protocol for optimal DNA recovery. The primers used can have a large effect on monitoring of environmental communities (55), so the use of probe-based assays could improve the results.

In the current study the sampling sites went through changes during the course of the experiment, the initial distinct organs such as the eyes, ears, and nose were no longer observable at later time points and samples were recovered from skeletal remains with minimal tissue mixed with maggot masses, and this new maggot-rich environment may have an effect on the local microbiome. Internal organs that were samples in other studies of human cadavers would also disappear as decomposition progresses (51). These factors suggest that reliable taphonomic use of microbial analysis may have to be restricted to early decomposition to ensure that the microbes that are analysed show consistency across the specimens, timeframes for post-mortem microbial analysis range from 10 – 20 days (51). At later stages of decomposition when environmental microbes are more influential there will be more site-specific effects that could prevent accurate extrapolation of data for PMI determinations. Soil microbial communities are affected by the nutrients leaching from cadavers and changes in soil microbial diversity could be more reliable indicators of PMI at later time points since the sampling sites will not decompose in the same manner as the actual cadaver (56, 57). In terms of sampling the greatest variation in bacterial abundance was found on the skin surface samples (T1 and T2) where sample recovery was not possible at 40-days (Figure 5). This coincided with a period of low rainfall and low temperature which may have dried the skin sufficiently to reduce bacterial population density on the surface of the animals further indicating that environment and state of decomposition can have a strong impact on microbial analysis of remains. It is essential to pair any microbial data with taphonomic observations to ensure accurate interpretation of any microbial data.

## Acknowledgements

This study was supported by technical staff from the School of Science, Western Sydney University.

## Author Contributions

HG, COM, SJ, conceived and designed the study. RL, BE conducted the experiments and data analysis. HG, COM, SJ contributed to the preparation of the manuscript.

## Availability of data and materials

The datasets used and/or analysed during the present study are available from the corresponding author on reasonable request.

## References

1. Houck MM. CSI: reality. Sci Am. 2006;295(1):84–9.

2. Metcalf JL, Wegener Parfrey L, Gonzalez A, Lauber CL, Knights D, Ackermann G, et al. A microbial clock provides an accurate estimate of the postmortem interval in a mouse model system. Elife. 2013;2:e01104.

3. Megyesi MS, Nawrocki SP, Haskell NH. Using accumulated degree-days to estimate the postmortem interval from decomposed human remains. J Forensic Sci. 2005;50(3):618–26.

4. Michaud JP, Moreau G. A statistical approach based on accumulated degree-days to predict decomposition-related processes in forensic studies. J Forensic Sci. 2011;56(1):229–32.

5. Campobasso CP, Di Vella G, Introna F. Factors affecting decomposition and Diptera colonization. Forensic Sci Int. 2001;120(1-2):18–27.

6. Goff ML. Early postmortem changes and stages of decomposition. Current concepts in forensic entomology. Netherlands: Springer 2009. p. 1–24.

7. Vass AA. The elusive universal post-mortem interval formula. Forensic Sci Int. 2011;204(1-3):34–40.

8. Marhoff-Beard SJ, Forbes SL, Green H. The validation of ’universal’ PMI methods for the estimation of time since death in temperate Australian climates. Forensic Sci Int. 2018;291:158–66.

9. Grivas CR, Komar DA. Kumho, Daubert, and the nature of scientific inquiry: implications for forensic anthropology. J Forensic Sci. 2008;53(4):771–6.

10. Munoz JI, Suarez-Penaranda JM, Otero XL, Rodriguez-Calvo MS, Costas E, Miguens X, et al. A new perspective in the estimation of postmortem interval (PMI) based on vitreous. J Forensic Sci. 2001;46(2):209–14.

11. Wallman JF. Winged evidence: forensic identification of blowflies. Australian Journal of Forensic Sciences. 2002;34(2):73–9.

12. Hayman J, Oxenham M. Estimation of the time since death in decomposed bodies found in Australian conditions. Australian Journal of Forensic Sciences. 2017;49(1):31–44.

13. Pechal JL, Crippen TL, Benbow ME, Tarone AM, Dowd S, Tomberlin JK. The potential use of bacterial community succession in forensics as described by high throughput metagenomic sequencing. Int J Legal Med. 2014;128(1):193–205.

14. Metcalf JL. Estimating the postmortem interval using microbes: Knowledge gaps and a path to technology adoption. Forensic Sci Int Genet. 2019;38:211–8.

15. Metcalf JL, Xu ZZ, Bouslimani A, Dorrestein P, Carter DO, Knight R. Microbiome Tools for Forensic Science. Trends Biotechnol. 2017;35(9):814–23.

16. Carter DO, Metcalf JL, Bibat A, Knight R. Seasonal variation of postmortem microbial communities. Forensic Sci Med Pathol. 2015;11(2):202–7.

17. Moitas B, Caldas IM, Sampaio-Maia B. Microbiology and postmortem interval: a systematic review. Forensic Sci Med Pathol. 2024;20(2):696–715.

18. Damann FE, Williams DE, Layton AC. Potential Use of Bacterial Community Succession in Decaying Human Bone for Estimating Postmortem Interval. J Forensic Sci. 2015;60(4):844–50.

19. Wang Z, Zhang F, Wang L, Yuan H, Guan D, Zhao R. Advances in artificial intelligence-based microbiome for PMI estimation. Front Microbiol. 2022;13:1034051.

20. Johnson HR, Trinidad DD, Guzman S, Khan Z, Parziale JV, DeBruyn JM, et al. A Machine Learning Approach for Using the Postmortem Skin Microbiome to Estimate the Postmortem Interval. PLoS One. 2016;11(12):e0167370.

21. Belk A, Xu ZZ, Carter DO, Lynne A, Bucheli S, Knight R, et al. Microbiome Data Accurately Predicts the Postmortem Interval Using Random Forest Regression Models. Genes (Basel). 2018;9(2).

22. Cruciani M, White PL, Barnes RA, Loeffler J, Donnelly JP, Rogers TR, et al. An Overview of Systematic Reviews of Polymerase Chain Reaction (PCR) for the Diagnosis of Invasive Aspergillosis in Immunocompromised People: A Report of the Fungal PCR Initiative (FPCRI)-An ISHAM Working Group. J Fungi (Basel). 2023;9(10).

23. Sezgin E, Terlemez G, Bozkurt B, Bengi G, Akpinar H, Buyuktorun I. Quantitative real-time PCR analysis of bacterial biomarkers enable fast and accurate monitoring in inflammatory bowel disease. PeerJ. 2022;10:e14217.

24. Ranjard L, Poly F, Nazaret S. Monitoring complex bacterial communities using culture-independent molecular techniques: application to soil environment. Res Microbiol. 2000;151(3):167–77.

25. Ott SJ, Musfeldt M, Ullmann U, Hampe J, Schreiber S. Quantification of intestinal bacterial populations by real-time PCR with a universal primer set and minor groove binder probes: a global approach to the enteric flora. J Clin Microbiol. 2004;42(6):2566–72.

26. Hauther KA, Cobaugh KL, Jantz LM, Sparer TE, DeBruyn JM. Estimating Time Since Death from Postmortem Human Gut Microbial Communities. J Forensic Sci. 2015;60(5):1234–40.

27. Galloway A, Birkby WH, Jones AM, Henry TE, Parks BO. Decay rates of human remains in an arid environment. J Forensic Sci. 1989;34(3):607–16.

28. Janaway RC, Percival SL, Wilson AS. Decomposition of human remains. In Microbiology and aging: Humana Press; 2009.

29. Marhoff SJ, Fahey P, Forbes SL, Green H. Estimating post-mortem interval using accumulated degree-days and a degree of decomposition index in Australia: a validation study. Australian Journal of Forensic Sciences. 2016;48(1):24–36.

30. Fierer N, Jackson JA, Vilgalys R, Jackson RB. Assessment of soil microbial community structure by use of taxon-specific quantitative PCR assays. Appl Environ Microbiol. 2005;71(7):4117–20.

31. Yang YW, Chen MK, Yang BY, Huang XJ, Zhang XR, He LQ, et al. Use of 16S rRNA Gene-Targeted Group-Specific Primers for Real-Time PCR Analysis of Predominant Bacteria in Mouse Feces. Appl Environ Microbiol. 2015;81(19):6749–56.

32. Blackwood CB, Oaks A, Buyer JS. Phylum- and class-specific PCR primers for general microbial community analysis. Appl Environ Microbiol. 2005;71(10):6193–8.

33. Faldynova M, Videnska P, Havlickova H, Sisak F, Juricova H, Babak V, et al. Prevalence of antibiotic resistance genes in faecal samples from cattle, pigs and poultry. Veterinarni Medicina. 2013;58(6):298–304.

34. Castillo M, Martin-Orue SM, Manzanilla EG, Badiola I, Martin M, Gasa J. Quantification of total bacteria, enterobacteria and lactobacilli populations in pig digesta by real-time PCR. Vet Microbiol. 2006;114(1-2):165–70.

35. Barman M, Unold D, Shifley K, Amir E, Hung K, Bos N, et al. Enteric salmonellosis disrupts the microbial ecology of the murine gastrointestinal tract. Infect Immun. 2008;76(3):907–15.

36. Rodrigues DF, da CJE, Ayala-Del-Rio HL, Pellizari VH, Gilichinsky D, Sepulveda-Torres L, et al. Biogeography of two cold-adapted genera: Psychrobacter and Exiguobacterium. ISME J. 2009;3(6):658–65.

37. Morton CO, Varga JJ, Hornbach A, Mezger M, Sennefelder H, Kneitz S, et al. The temporal dynamics of differential gene expression in Aspergillus fumigatus interacting with human immature dendritic cells in vitro. PLoS One. 2011;6(1):e16016.

38. Damron FH, Owings JP, Okkotsu Y, Varga JJ, Schurr JR, Goldberg JB, et al. Analysis of the Pseudomonas aeruginosa regulon controlled by the sensor kinase KinB and sigma factor RpoN. J Bacteriol. 2012;194(6):1317–30.

39. Forger LV, Woolf MS, Simmons TL, Swall JL, Singh B. A eukaryotic community succession based method for postmortem interval (PMI) estimation of decomposing porcine remains. Forensic Sci Int. 2019;302:109838.

40. Forbes SL, Perrault KA, Stefanuto PH, Nizio KD, Focant JF. Comparison of the decomposition VOC profile during winter and summer in a moist, mid-latitude (Cfb) climate. PLoS One. 2014;9(11):e113681.

41. Meyer J, Anderson B, Carter DO. Seasonal variation of carcass decomposition and gravesoil chemistry in a cold (Dfa) climate. J Forensic Sci. 2013;58(5):1175–82.

42. Archer MS. Rainfall and temperature effects on the decomposition rate of exposed neonatal remains. Sci Justice. 2004;44(1):35–41.

43. Lloyd-Price J, Abu-Ali G, Huttenhower C. The healthy human microbiome. Genome Med. 2016;8(1):51.

44. Preiswerk D, Walser JC, Ebert D. Temporal dynamics of microbiota before and after host death. ISME J. 2018;12(8):2076–85.

45. DeBruyn JM, Hauther KA. Postmortem succession of gut microbial communities in deceased human subjects. PeerJ. 2017;5:e3437.

46. Pechal JL, Schmidt CJ, Jordan HR, Benbow ME. A large-scale survey of the postmortem human microbiome, and its potential to provide insight into the living health condition. Sci Rep. 2018;8(1):5724.

47. Hyde ER, Haarmann DP, Petrosino JF, Lynne AM, Bucheli SR. Initial insights into bacterial succession during human decomposition. Int J Legal Med. 2015;129(3):661–71.

48. Benbow ME, Pechal JL, Lang JM, Erb R, Wallace JR. The Potential of High-throughput Metagenomic Sequencing of Aquatic Bacterial Communities to Estimate the Postmortem Submersion Interval. J Forensic Sci. 2015;60(6):1500–10.

49. Olakanye AO, Thompson T, Ralebitso-Senior TK. Shifts in soil biodiversity-A forensic comparison between Sus scrofa domesticus and vegetation decomposition. Sci Justice. 2015;55(6):402–7.

50. Jo YJ, Tagele SB, Pham HQ, Jung Y, Ibal JC, Choi S, et al. In Situ Profiling of the Three Dominant Phyla Within the Human Gut Using TaqMan PCR for Pre-Hospital Diagnosis of Gut Dysbiosis. Int J Mol Sci. 2020;21(6).

51. Javan GT, Finley SJ, Smith T, Miller J, Wilkinson JE. Cadaver Thanatomicrobiome Signatures: The Ubiquitous Nature of Clostridium Species in Human Decomposition. Front Microbiol. 2017;8:2096.

52. Deschaght P, Janssens M, Vaneechoutte M, Wauters G. Psychrobacter isolates of human origin, other than Psychrobacter phenylpyruvicus, are predominantly Psychrobacter faecalis and Psychrobacter pulmonis, with emended description of P. faecalis. Int J Syst Evol Microbiol. 2012;62(Pt 3):671–4.

53. Zhao W, Wang Y, Liu S, Huang J, Zhai Z, He C, et al. The dynamic distribution of porcine microbiota across different ages and gastrointestinal tract segments. PLoS One. 2015;10(2):e0117441.

54. Iancu L, Bonicelli A, Procopio N. Decomposition in an extreme cold environment and associated microbiome-prediction model implications for the postmortem interval estimation. Front Microbiol. 2024;15:1392716.

55. Fredriksson NJ, Hermansson M, Wilen BM. The choice of PCR primers has great impact on assessments of bacterial community diversity and dynamics in a wastewater treatment plant. PLoS One. 2013;8(10):e76431.

56. Singh B, Minick KJ, Strickland MS, Wickings KG, Crippen TL, Tarone AM, et al. Temporal and Spatial Impact of Human Cadaver Decomposition on Soil Bacterial and Arthropod Community Structure and Function. Front Microbiol. 2017;8:2616.

57. Cobaugh KL, Schaeffer SM, DeBruyn JM. Functional and Structural Succession of Soil Microbial Communities below Decomposing Human Cadavers. PLoS One. 2015;10(6):e0130201.

